# Microbiome-based environmental monitoring of a dairy processing facility highlights the challenges associated with low microbial-load samples

**DOI:** 10.1101/2020.09.11.292862

**Authors:** Aoife J. McHugh, Min Yap, Fiona Crispie, Conor Feehily, Colin Hill, Paul D. Cotter

**Affiliations:** Food Bioscience Department, Teagasc Food Research Centre, Cork, Ireland; School of Microbiology, University College Cork, Cork, Ireland; APC Microbiome Ireland, Cork, Ireland

## Abstract

Food processing environments can harbor microorganisms responsible for food spoilage or foodborne disease. Efficient and accurate identification of microorganisms throughout the food chain can allow the identification of sources of contamination and the timely implementation of control measures. Currently, microbial monitoring of the food chain relies heavily on culture-based techniques. These assays are determined on the microbes expected to be present in the environment, and thus do not cater for unexpected contaminants. Many culture-based assays are also unable to distinguish between undesirable taxa and closely related harmless species. Furthermore, even when multiple culture-based approaches are used in parallel, it is still not possible to comprehensively characterize the entire microbiology of a food-chain sample.

High throughput DNA sequencing represents a potential means through which microbial monitoring of the food chain can be enhanced. While sequencing platforms, such as the Illumina MiSeq, NextSeq and NovaSeq, are most typically found in research or commercial sequencing laboratories, newer portable platforms, such as the Oxford Nanopore Technologies (ONT) MinION, offer the potential for rapid analysis of food chain microbiomes. In this study, having initially assessed the ability of rapid MinION-based sequencing to discriminate between different microbes within a simple mock metagenomic mixture of related food spoilage, spore-forming microorganisms. Subsequently, we proceeded to compare the performance of both ONT and Illumina sequencing for environmental monitoring of an active food processing facility.

Overall, ONT MinION sequencing provided accurate classification to species level, which was comparable to Illumina-derived outputs. However, while the MinION-based approach provided a means of easy library preparations and portability, the high concentrations of DNA needed to run the rapid sequencing protocols was a limiting factor, requiring the random amplification of template DNA in order to generate sufficient material for analysis.

## Introduction

Dairy processing environments harbor microorganisms that have the potential to contaminate food before and during processing (Gleeson, O&Connell and Jordan, 2013; Doyle *et al*., 2017; Faille *et al*., 2014; Wang *et al*., 2019; Fysun *et al*., 2019). Some of these microorganisms have the potential to cause spoilage or be pathogenic (Doyle *et al*., 2015; Cho *et al*., 2018; Sadiq *et al*., 2016; Burgess, Lindsay and Flint, 2010). Routine environmental monitoring is carried out in food processing environments for this reason, and usually involves the use of swabbing and agar plating to determine total numbers of general (e.g., total bacteria count) or specific (generally potentially spoilage-associated or pathogenic species) categories of microorganisms (Cho *et al*., 2018). These analyses frequently involve phenotype-based agar assays, some of which can yield high false positive rates (Doyle, O&Toole and Cotter, 2018; Tallent *et al*., 2012). These approaches are further limited by the fact that they do not provide information about non-targeted species or indeed the microbial population as a whole.

DNA sequencing methods have recently been applied to dairy and environmental samples to determine the microbial population composition and enable source tracking (Doyle *et al*., 2017; McHugh *et al*., 2018; Fretin *et al*., 2018; Cho *et al*., 2018, McHugh *et al*., 2020). High throughput metagenomic sequencing can provide greater insights into the taxonomic composition of populations present in these environments than culture based methods. Specifically it uncovers information relating to the functional potential of species and strains present, including virulence and spoilage properties. Despite these benefits, high throughput metagenomic sequencing approaches typically require expensive reagents and platforms as well as personnel skilled in molecular biology, data generation and interpretation. These requirements limit their routine implementation in manufacturing facilities. Some of these issues have the potential to be addressed through use of portable DNA sequencing devices such as the Oxford Nanopore Technologies (ONT) MinION. The MinION’s portability and work flows are designed to facilitate their use by less experienced personnel and could allow easier detection and identification of the causative agents of microbial contamination. Such approaches have recently been tested in a clinical setting to identify causative agents of disease from metagenomic samples (Charalampous *et al*., 2019), including studies where the results were compared with those generated through Illumina sequencing (Quick *et al*., 2017; Kafetzopoulou *et al*., 2018) or culture-based analysis (Sanderson *et al*., 2018). This approach has yet to be applied to food processing settings for environmental monitoring.

As a proof-of-concept, we conducted a study to determine the ability of MinION-based rapid sequencing to correctly classify a simple, four strain, mock community of highly related spore-forming microorganisms of relevance to the dairy processing chain. Prompted by this initial analysis, we proceeded to compare the outputs of MinION-based rapid sequencing to Illumina-based, and culture-based methods to characterize the microbiota of environmental swabs collected from a food processing facility. Overall, MinION-based approaches were comparable to the Illumina sequencing equivalent in terms of species level taxonomic classification. However, the requirement of high concentration and quality input DNA for the routine implementation of MinION sequencing was a limitation due to the environment tested. To overcome this, random amplification of template DNA was required. Regardless, the potential benefits of the routine application of metagenomic sequencing to food processing environments were clear.

## Results

### MinION sequencing accurately identified, and distinguished between, genomic DNA from four related, dairy environment-associated, sporeformers

Metagenomic DNA representing a simple mock community of 4 related dairy processing-associated, spore-forming contaminants, i.e., *Bacillus cereus, Bacillus. thuringiensis, Bacillus licheniformis, Geobacillus stearothermophilus*, was sequenced using ONT MinION rapid sequencing kits. This proof-of-concept exercise was performed to determine the extent to which MinION-based sequencing could identify, and discriminate between, related, and in some cases difficult to distinguish, microorganisms found in dairy processing environments. Amplicon 16S rRNA-based sequencing of the simple mock metagenomic DNA using the ONT 16S barcoding kit SQK-RAB204 resulted in 996,441 reads following rebasecalling by albacore. These reads contained a total of 1,454,835,092 bases with an average read length of 1460 bp and a median read length of 1561 bp. 16S rRNA reads aligned by BLASTn to the Silva 16S database (version 132) with MEGAN 6 classification resulted in successful identification of three out of the 4 species. The fourth strain, *G. stearothermophilus* DSM 458, was correctly identified to the genus level only (Figure 1A).

**Figure 1.**
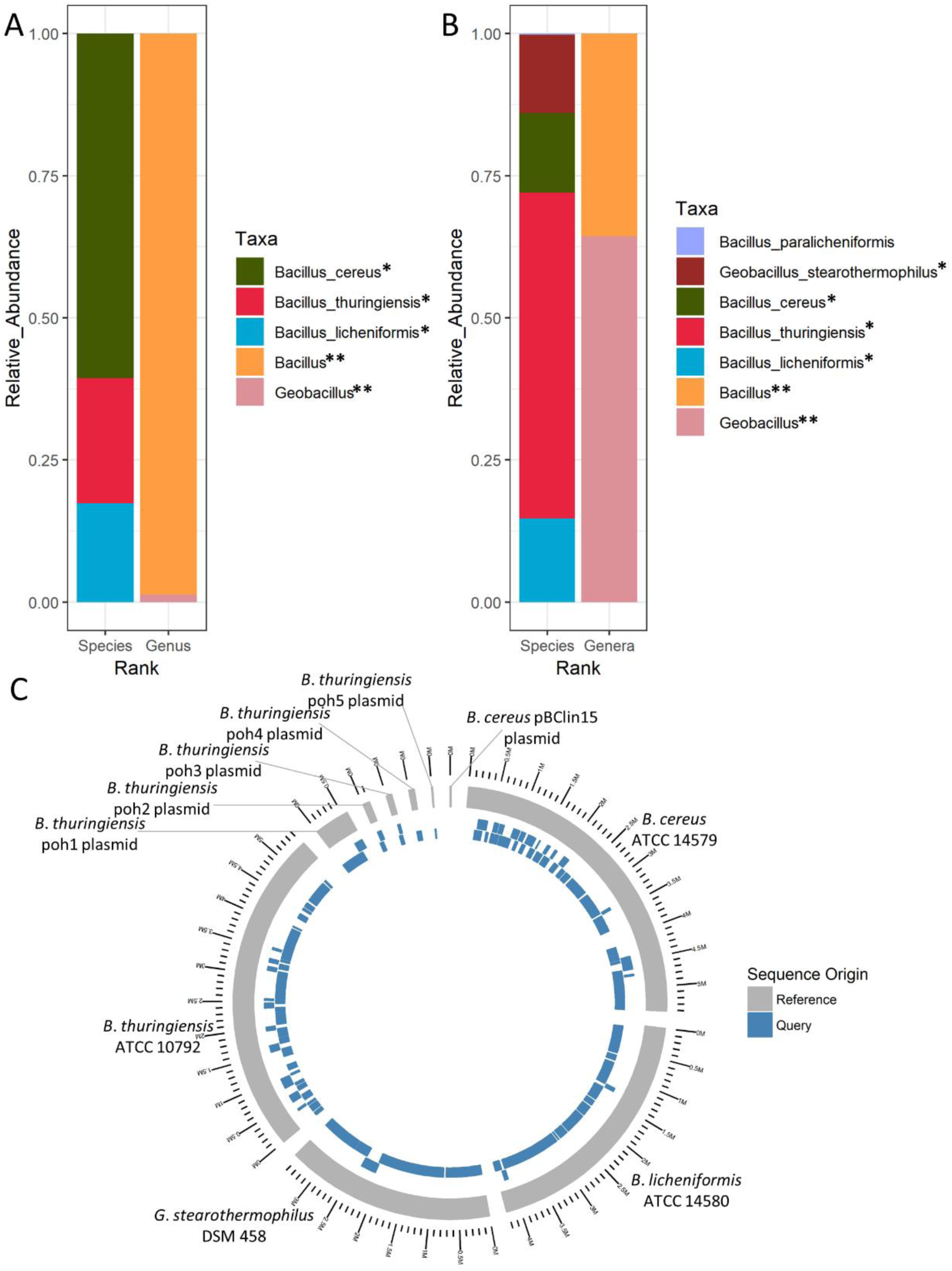
Mock community analysis. MinION sequencing followed by MEGAN taxonomic classification of a simple mock community. A. Taxonomic classification following 16S sequencing. Expected species are denoted with *, while expected genera are denoted **. B. Taxonomic classification following rapid WMGS. Expected species are denoted with *, while expected genera are denoted **. C. *De novo* assembly of genomes by the canu assembler, followed by mapping back to original known genomes, to illustrate coverage at 97% identity. 4 genomes, with 6 plasmids illustrated, of which 4 genomes and 5 plasmids had sequences aligned at 97% identity.

Rapid whole metagenome sequencing (WMGS) of the mock community using the SQK-RAD004 kit resulted in 97,503 reads following rebasecalling by albacore and adaptor removal. These 97,503 reads contained a total of 750,359,905 bases with an average read length of 7696 bases and a median of 5762 bases. LAST alignment against the nr database followed by MEGAN long read (LR) lowest common ancestor (LCA) analysis resulted in 74.76% bases being classified to some taxonomic level. Of these, 42.63% were classified to species level, 46.28% classified to species group level and 8.15% classified to genus level, accounting for 97.06% of classified reads. 64.37% of bases classified to genus level only were attributed to *Geobacillus*, with the remaining 35.63% classified as *Bacillus* (Figure 1B). Of the sequences classified to the species level, 57.26% of bases were attributed to *Bacillus thuringiensis*, 14.74% were attributed to *Bacillus licheniformis*, 13.98% were attributed to *Bacillus cereus*, 13.8% were attributed to *Geobacillus stearothermophilus* and, 0.21% misassigned as *Bacillus paralicheniformis* (Figure 1B). *De novo* assembly of raw reads from the rapid sequencing reads using the canu (version 1.7) assembler (Koren *et al*., 2017) resulted in 104 contigs and mapping back of reads to references resulted in good coverage up to 97% identity (Figure 1C). The 4 reference strains genomes included 6 plasmids, corresponding to 10 contiguous stretches of DNA. Nine of these 10 contigs were identified following sequence assembly, the exception being pBClin15, a 15 kb plasmid from *B. cereus* (Figure 1C). 99.59% of the assembled bases aligned to the reference genomes and, of the reference genomes, 98.27% aligned to the assembled MinION sequences (Supplemental Table 1).

**Table 1.** High quality MAGs. Taxonomy was assigned to metaBAT2 binned contigs by Megan LR and open reading frames of these contigs by Kaiju. If more than one species assigned, the bold species represents the top hit per classifier. Bin quality determined by checkM.

| Month | Sample | Bin | Kaiju assignment | Megan assignment | Percent Complete | Percent Contamination |
| --- | --- | --- | --- | --- | --- | --- |
| October | Positive control (mock) | 2 | *Geobacillus stearothermophilus* | *Geobacillus stearothermophilus* | 91.37 | 1.1235 |
| October | Evaporator Drain | 5 | ***Planococcus plakortidis***/<br>*Planococcus maitriensis*/<br>*Planococcus maritimus* | ***Planococcus plakortidis***/<br>*Planococcus maritimus*/<br>*Planococcus rifietoensis* | 82.41 | 0.6622 |
| October | Gasket/Flow Plate Seal | 5 | ***Exiguobacterium acetylicum***/<br>*Exiguobacterium sp. RIT341*/<br>*Exiguobacterium indicum* | ***Exiguobacterium indicum***/<br>*Exiguobacterium acetylicum* | 81.9 | 0.6578 |
| October | Table | 14 | *Enterococcus casseliflavus* | *Enterococcus casseliflavus* | 82.77 | 1.4622 |
| November | Positive control (mock) | 1 | *Bacillus licheniformis* | *Bacillus licheniformis* | 94.23 | 0.4149 |
| November | Evaporator Drain | 7 | *Paracoccus chinensis* | *Paracoccus chinensis* | 93.03 | 1.1235 |
| November | Gasket/Flow Plate Seal | 1 | *Macrococcus caseolyticus* | *Macrococcus caseolyticus* | 90.35 | 1.1049 |
| November | External Dryer Balance Tank | 3 | *Nesterenkonia massiliensis* | *Nesterenkonia massiliensis* | 91.32 | 1.07 |
| December | Positive control (mock) | 3 | *Bacillus licheniformis* | *Bacillus licheniformis* | 96.29 | 0.0829 |
| December | External Dryer Balance Tank | 1 | ***Kocuria sp. CPCC 104605***/<br>*Kocuria sp. ZOR0020* | *Kocuria sp. ZOR0020* | 81.35 | 0.6578 |

### Shotgun sequencing of environmental dairy processing samples through MinION and NextSeq sequencing provided comparable taxonomic classifications

Prompted by the successful use of MinION-based sequencing to characterise the mock metagenomic community DNA, the technology was applied to study the microbiota of a food processing facility and to compare outputs with those derived through NextSeq (Illumina)-based sequencing. Eight locations in a single processing facility were swabbed on three different days across October, November, and December 2018, each after cleaning in place (CIP) but before the next round of dairy processing (Figure 2). These eight locations comprised a table, door, wall, gaskets/flow plate seals, external surface of dryer balance tank, internal surface of dryer balance tank, external surface of evaporator, and drain beside evaporator. These swabs were prepared for sequencing, along with a series of negative controls and a positive control, consisting of the simple mock metagenomic community used previously. For MinION sequencing, rapid sequencing of multiple displacement amplification (MDA)-generated template DNA from 36 samples, used to address the relatively high quantities of DNA required for library preparation, was carried out using the SQK-RBK004 rapid barcoding sequencing kit. After processing, a total of 899,306 reads were generated, containing a total of 1,648,724,928 bases with an average read length of 1,833 bases and median of 926 bases per read (and an average of 45,797,915 bases and 24,980.7 reads per sample). LAST alignment against the nr database followed by MEGAN long read (LR) lowest common ancestor (LCA) analysis resulted in 62% of bases being classified to some taxonomic level. Of these, 29.11% were classified to species level and 38.36% classified to genus level, accounting for 67.47% of classified reads. A total of 59 species were detected at > 5% relative abundance in at least one sample by MEGAN (Supplemental Figure 2).

**Figure 2.**
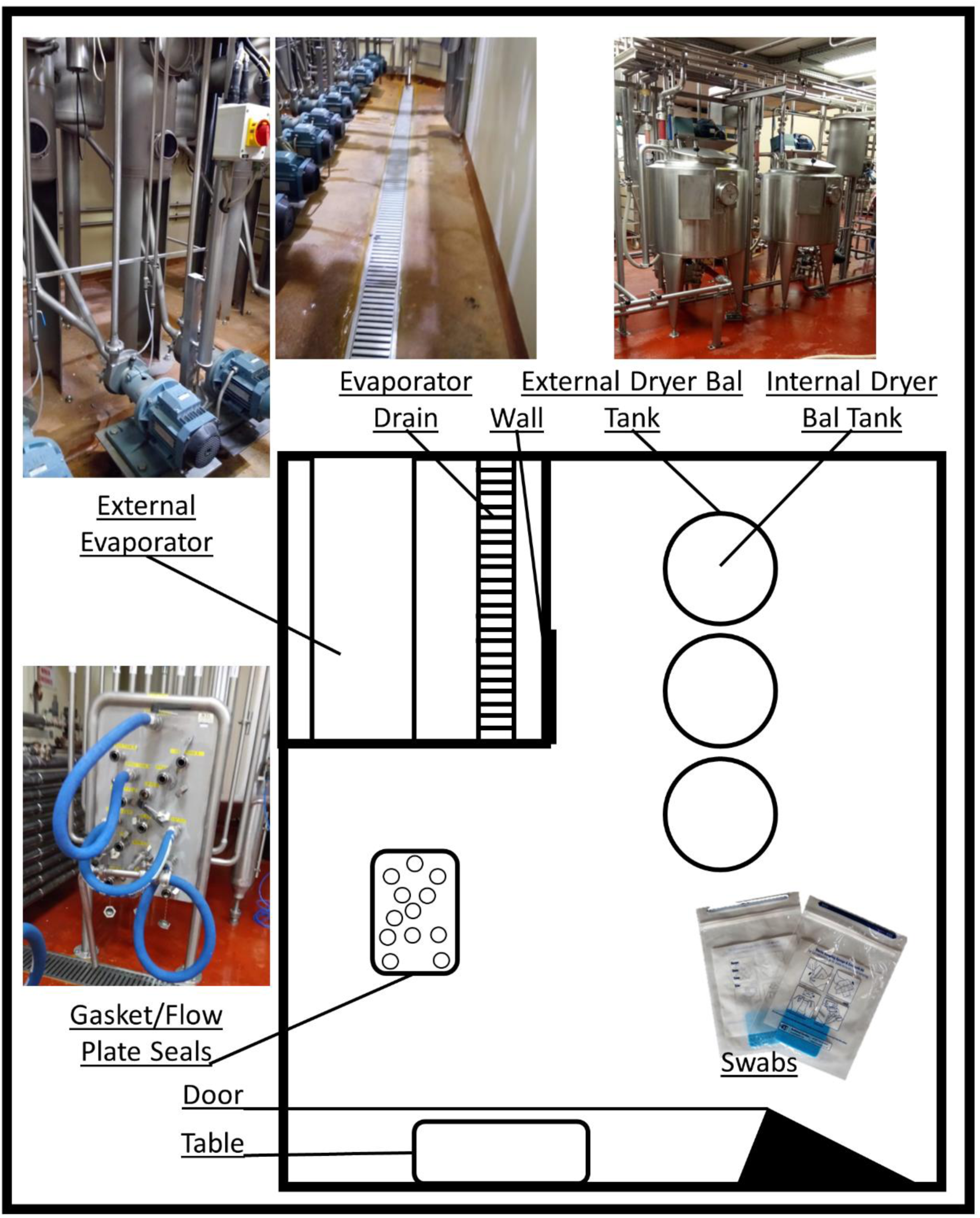
Schematic of dairy processing facility sampling areas. Dairy processing facility schematic includes the 8 areas sampled in each of October, November, and December 2018. Areas were sampled post CIP and prior to the recommencement of processing.

Other shotgun sequencing-based approaches were employed to study the microbiomes of these environmental samples for comparative purposes. These included Illumina-based sequencing of MDA and non-MDA DNA, as well as of metagenomic DNA extracted from easily cultured metagenomic DNA to allow a comparison with the species that grow when traditional culturing-based approaches are employed. This Illumina (NextSeq)-based sequencing of 93 samples produced 734,909,370 reads containing 150 bases each with an average of 7,902,251 reads per sample. To allow a comparison with MinION outputs, and to avoid discrepancies through use of different bioinformatic pipelines, Diamond alignment against the nr database followed by MEGAN 6 lowest common ancestor (LCA) analysis was employed and resulted in 78% reads being classified to some taxonomic level. Of these, 10.8% were classified to the species level and 39.6% classified to the genus level, accounting for 50.3 % of classified bases. In comparison, Kraken2 and Bracken classification resulted in 61% reads classified to some taxonomic level, with 99% of those classified being classified to species level. This approach did not correctly classify the composition of the mock community (positive control) (Supplemental Figure 3). Similarly, MetaPhlAn2 did not correctly classify all of the species of the mock community (Supplemental Figure 4), with both classifiers incorrectly classifying at least one species. Interestingly, both classifiers misclassified different species, whereby Bracken misclassified *B. licheniformis* as a *Bacillus* phage, and MetaPhlAn2 did not differentiate between *B. cereus* and *B. thuringiensis*. Additionally, MetaPhlAn2 only classified the *G. stearothermophilus* to genus level.

Using the MEGAN classification, which correctly classified the simple mock community, 108 species were identified at > 5% relative abundance in at least one sample from all MinION and NextSeq sequenced samples (Figure 3). Species level classification by MEGAN revealed consistencies between corresponding NextSeq-and MinION-sequenced samples (Figure 3). Overall, reads corresponding to *Kocuria sp*. WRN011 were detected at the highest relative abundance. This taxon was detected in multiple locations, at each time-point, in both the MinION, and corresponding NextSeq, MDA-generated samples. Its relative abundance was highest in the evaporator drain samples at each time point. *Kocuria sp*. ZOR0020 was present in high relative abundance in external dryer balance tank swabs in both MinION-and NextSeq-MDA sequenced MDA samples (Figure 3). Other dominant species included *Acinetobacter johnsonnii* in gasket/flow plate seals (MinION and Illumina), *Micrococcus luteus* in evaporator drain (MinION and Illumina sequenced samples), *Enterococcus faecium* from the inside of the dryer balance tank as well as many other October and November samples (MinION and MDA amplified Illumina sequencing), *Klebsiella pneumonia* in many December samples regardless of sequencing approach and *Enterococcus casseliflavus* in many samples from October and November (high relative abundance in MinION sequenced samples and at lower abundance in the corresponding MDA Illumina sequenced samples) (Figure 3). *Exiguobacterium sibiricum* was also detected in high relative abundance in MinION sequenced October and November door samples. It was also at lower relative abundances in many other October and November samples and in the corresponding Illumina sequenced door samples.

**Figure 3.**
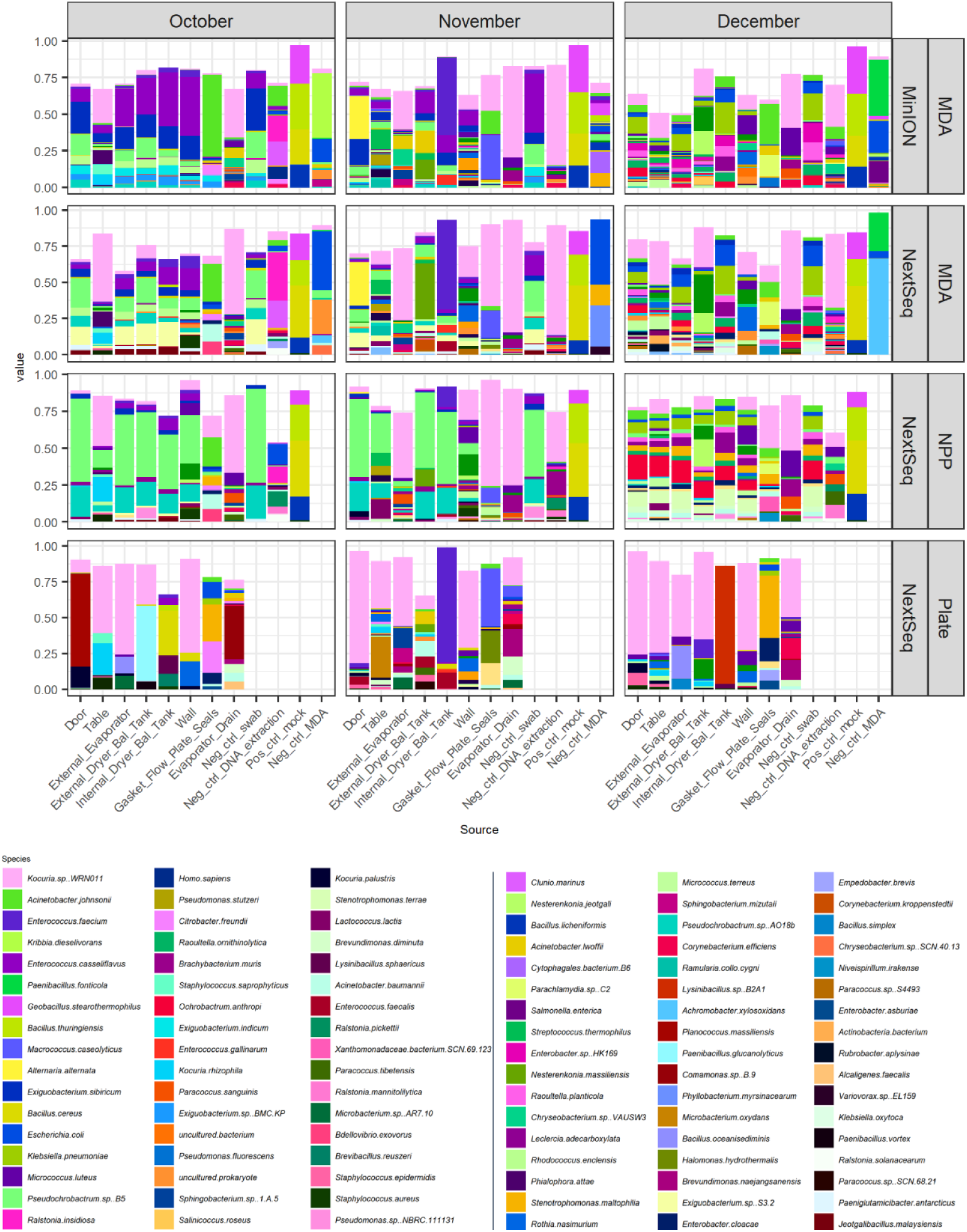
Species level classification of MinION and NextSeq sequenced environmental samples. Taxonomic assignment of MinION and NextSeq sequenced samples generated following the use of different pre-processing and sequencing methods. Pre-processing methods include MDA amplification, no pre-processing (NPP), and spread plating on BHI before washing colonies, pelleting, and treating as a metagenomic sample (Plate). Species level classification was performed using LAST (for MinION) and Diamond (for NextSeq) alignment of reads against the NR database and classification with MEGAN (LR for MinION). Species present in at least 5% in at least one sample are shown.

There were some notable sequencing platform-dependent differences.

*Exiguobacterium sp*. S3.2 and *Pseudochrobactrum sp* B5 were present at higher relative abundance in October and November MDA Illumina NextSeq sequences compared to MinION sequences and *Enterobacter sp*. HK169 was detected in December MinION samples, but not corresponding Illumina samples (Figure 3). Species level taxonomic identification was performed on negative controls also. Many species were specific to negative controls, including *Kribbia dieselivorans* and *Cytophagales bacterium* B6, detected at a high relative abundance in MinION sequenced MDA negative controls, and *Paenibacillus fonticola*, detected at high relative abundance in both MinION and Illumina sequenced MDA negative controls. There was also a high relative abundance of *Escherichia coli* in MDA negative controls, with *Salmonella enterica* in the December samples, in both MinION sequences and corresponding Illumina sequences. *Ralsonia insidiosa* was also seen above 0.2% exclusively in negative controls. However, there was some overlap with species identified in negative controls also identified in environmental samples. In particular, the swab negative control for both MDA MinION and MDA NextSeq from each month are similar to results generated from swabbing of the internal of the dryer balance tank, which are the environmental samples with the lowest environmental load (Supplemental Table 2). *Kocuria sp*., *Acinetobacter johnsonnii, Enterococcus casseliflavus, Klebsiella pneumoniae, Exiguobacterium sibricum, Enterococcus casseliflavus, Pseudochrobctrum sp* B5, *Enterobacter sp* HK169 and *Raoultella planticola* are all seen in negative controls (Figure 3). These findings highlight the risks of relying on data from samples will a low microbial load and the importance of including negative controls.

Metagenome-assembled genomes (MAGs) were extracted from assemblies of combined Illumina MDA and MinION MDA sequences. This resulted in 162 bins, of which 10 were high quality at > 80% complete and < 10% contamination (Table 1). 7 of the 10 MAGs were from environmental isolates, with 3 out of 10 being the positive control species used. From the remaining MAGs, 3 out of 7 environmental isolates could not be definitively assigned at the species level, being assigned as each of a number of species at similar levels of relative abundances. These MAGs were assigned at the genus level as *Planococcus, Exiguobacterium* and *Kocuria* and were sourced from the October evaporator drain, gasket/flow plate seal and external dryer balance tank, respectively. The MAGs that were assigned at the species level were an *Enterococcus casseliflavus* from the October table swab sample, a *Paracoccus chinensis* from the November evaporator drain, a *Micrococcus casseolyticus* from the November gasket/flow plate seal and a *Nesterenkonia massiliensis* from the November external of dryer balance tank sample (Table 1).

### MDA amplification introduced bias towards the detection of some species

In order to determine the potential for bias arising from MDA pre-processing, outputs from MDA-generated NextSeq sequencing were compared to non-MDA derived NextSeq (NPP). Higher relative abundances of *Pseudochrobactrum sp*. B5 and *Pseudochrobactrum sp*. AO18b were seen in October and November NPP samples compared to the MDA-amplified equivalents (Figure 3). Overall, the NPP samples we found to be less diverse than their MDA counterparts (Figure 4A).

**Figure 4.**
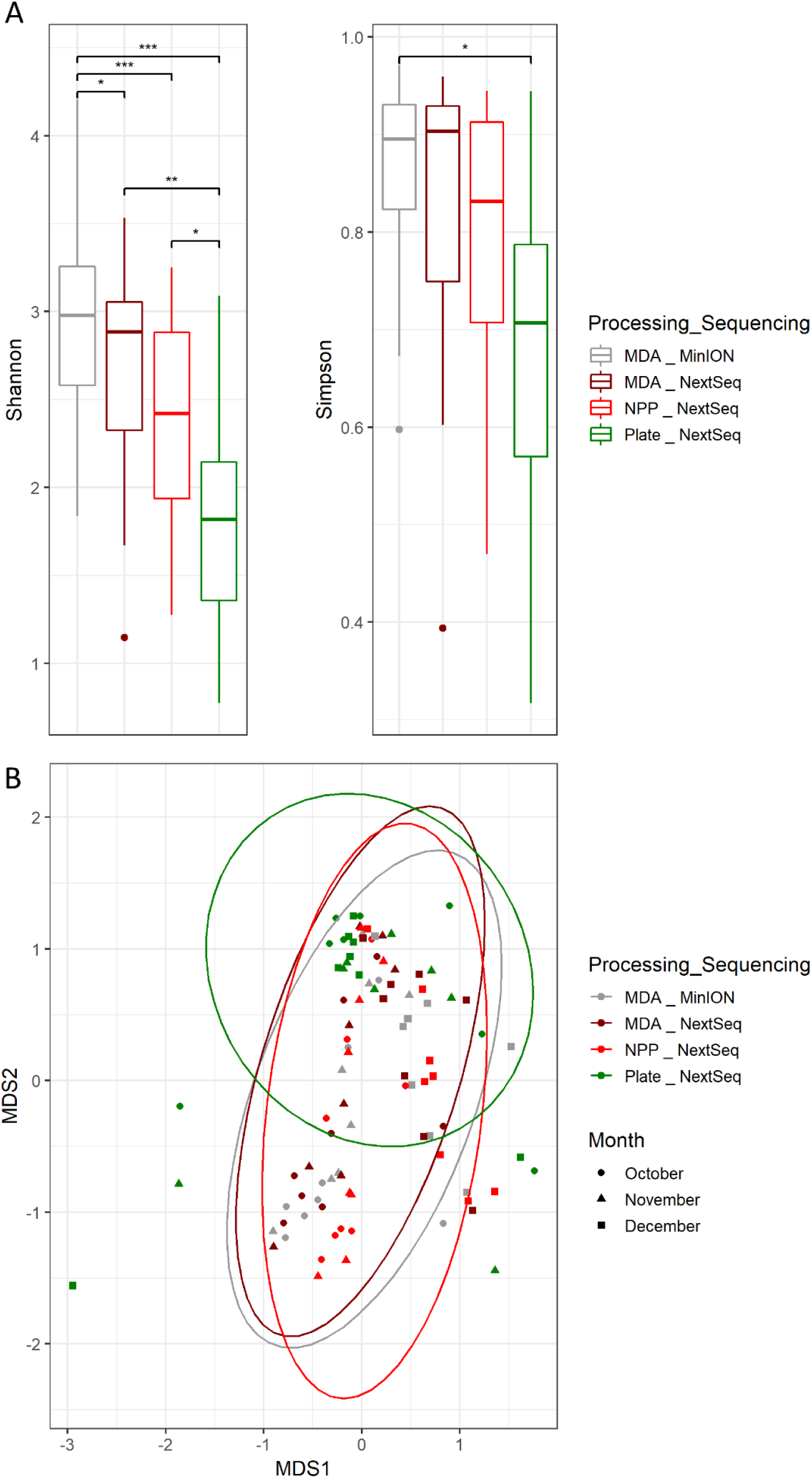
Diversity analysis. A. Shannon and Simpson alpha diversity analysis. B. Bray Curtis multiple displacement scaling (MDS) beta diversity analysis. (*** = p < 0.001, ** = p < 0.01, * = p < 0.05). Controls are excluded from these calculations and figures.

### Culture-based analyses introduced a selection bias

In order to determine to what extent culture-dependent and –independent approaches provided different outputs, a comparison between NPP NextSeq-generated sequences and those resulting from sequencing of pools of easily cultured colonies (Plate samples) was performed. Sequences generated from Plate samples were noted to be significantly less diverse (Figure 4A), however the Plate samples clustered with the non-cultured samples when beta-diversity was analysed (Figure 4B). A number of the species detected were similar to species identified in the corresponding non-cultured samples (NPP and MDA amplified). Overall, *Kocuria sp* WRN011, was detected in all samples in which it had previously been identified through culture-independent approaches. *Enterococcus faecium*, the species found at highest relative abundance in all internal dryer balance tank samples from November (i.e., MDA MinION, NextSeq MDA, NPP and Plate; Figure 3) was also detected. Pre-culturing enriched some species that had been identified at low relative abundance in metagenomic NPP and MDA samples. These included *Planococcus massiliensis* (October door sample), *Microbacterium oxydans* (November Table sample), *Acinetobacter baumannii* (November external dryer balance tank) and *Lysinibacillus sp* B2A1 (December internal dryer balance tank; Figure 3).

### Genus level classification highlighted further culture-based selection bias

As some genera could not be distinguished at species level, genus level assignments were also investigated and compared. MEGAN LCA analysis identified sequences that could not be more accurately classified to species level, and assigned these as far as genus level only. A combined 56 genera were identified between MinION, NextSeq (both at > 5% relative abundance) and Sanger sequencing. Fifteen of these 56 genera were identified in samples from all 3 sequencing types (Supplemental Figure 5). Sanger sequencing involved partial 16S rRNA sequencing of morphologically different colonies from BHI plates, including total spread plate (TBC), thermophilic enriched spore pasteurised (ST) and mesophilic enriched spore pasteurised (SM) tests (Supplemental Table 2; Figure 5). There was agreement between Sanger sequencing of isolates and next generation sequencing of plate samples with respect to *Kocuria, Acinetobacter* and *Lysinibacillus* (Figure 5). Some genera identified in Plate NextSeq samples and Sanger sequences had not been seen in high relative abundance in corresponding culture-independent NextSeq or MinION sequencing. These included *Microbacterium* in the November table sample and *Lysinibacillus* in the December internal dryer balance tank (Figure 5).

**Figure 5.**
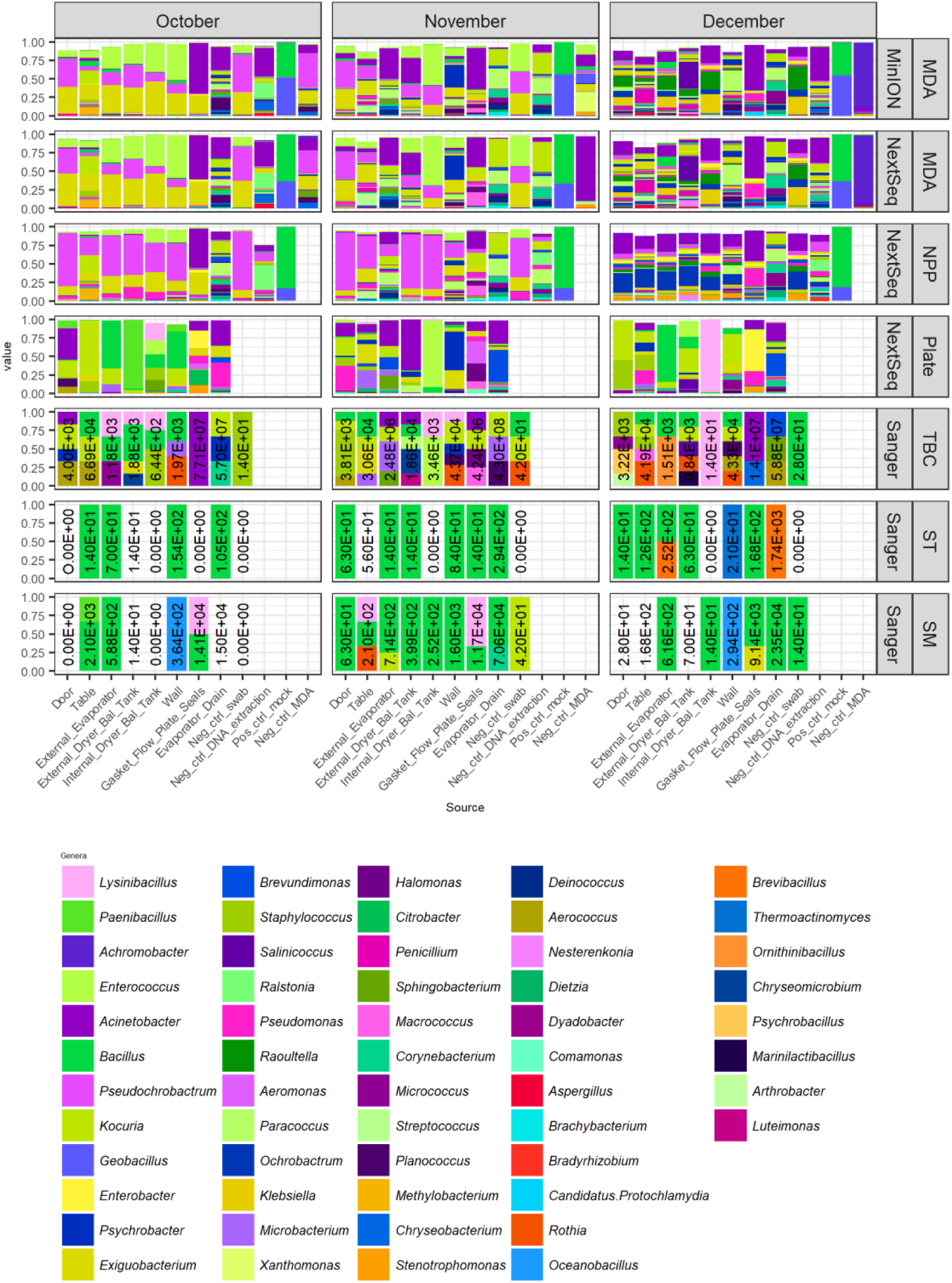
Genus level classification of environmental samples and controls following different pre-processing methods and sequencing methods. MEGAN LCA based genera level classification of MinION and NextSeq sequences. Also depicted are Sanger results to genus level for morphologically different colonies from each sample (TBC) along with thermophilic sporeformer enriched (ST) and mesophilic sporeformer enriched (SM) counts. Also included are CFU / swab counts for each culturing type. Sanger results represent relative abundance of a subset of morphologically distinct isolates rather than total isolates.

Overall, Sanger sequencing of 16S variable region of TBC isolates corresponded well with NextSeq ‘Plate’ sequencing but fewer genera were identified per sample. This may in part be due to only very morphologically distinct isolates being selected for Sanger sequencing. Counts per swab are also included. At all timepoints the gasket/flow plate seals and the evaporator drains had highest CFU / swab, with on average 3.18 × 10^7^ CFU / swab and 1.82 × 10^8^ CFU / swab each. These two areas also had the highest mesophilic spore count with an average of 1.17 × 10^4^ CFU / swab and 3.64 x10^4^ CFU / swab each (Figure 5, Supplemental Table 2).

### Relatively few significant differences in relative abundance of species and genus level taxonomic classification due to sequencing and pre-processing approaches

Overall, only 6 out of 108 species had significantly different relative abundance between environmental samples (excluding controls) due to sample processing or sequencing method, based on Pairwise Wilcoxon rank sums test using Benjamini Hochberg *p*-value correction analysis of sequential pairs (Supplemental Figure 6). *Enterococcus casseliflavus, Acinetobacter lwoffii* and *Acinetobacter johnsonii* had significantly higher relative abundance in MDA MinION sequenced samples than MDA NextSeq sequenced samples, whereas *Kocuria sp*. WRN011 was identified at significantly higher relative abundance in MDA NextSeq samples than MDA MinION samples. *Pseudochrobactrum sp* B5 was detected at significantly higher relative abundance in NPP NextSeq samples than MDA processed NextSeq samples, whereas *Exiguobacterium sibricum* was detected at significantly higher relative abundance in MDA NextSeq samples compared to NPP NextSeq samples (Figure 6A). Genera that had significantly different relative abundances, depending on whether MinION or NextSeq sequencing approaches were used, were also identified. In this case a greater number of significantly different taxa was observed, with 24 genera out of a total of 46 being significantly different as a consequence of the sample processing or sequencing method used (Figure 6B, Supplemental Figure 7). Six genera differed significantly between more than one pairwise group (Figure 6B). *Pseudochrobactrum* was present at significantly different relative abundances across all 3 pairwise groups (i.e., MDA MinION and MDA NextSeq, MDA NextSeq and NPP NextSeq, and NPP NextSeq and Plate NextSeq). *Exiguobacterium* and *Planococcus* were present at significantly different relative abundances between MDA MinION and MDA NextSeq as well as MDA NextSeq and NPP NextSeq. *Bacillus, Staphylococcus* and *Ochrobactrum* were present at significantly different relative abundances between MDA NextSeq and NPP NextSeq as well as NPP NextSeq and Plate NextSeq. The remaining 18 genera only differed across one pair of analyses (Figure 6B).

**Figure 6.**
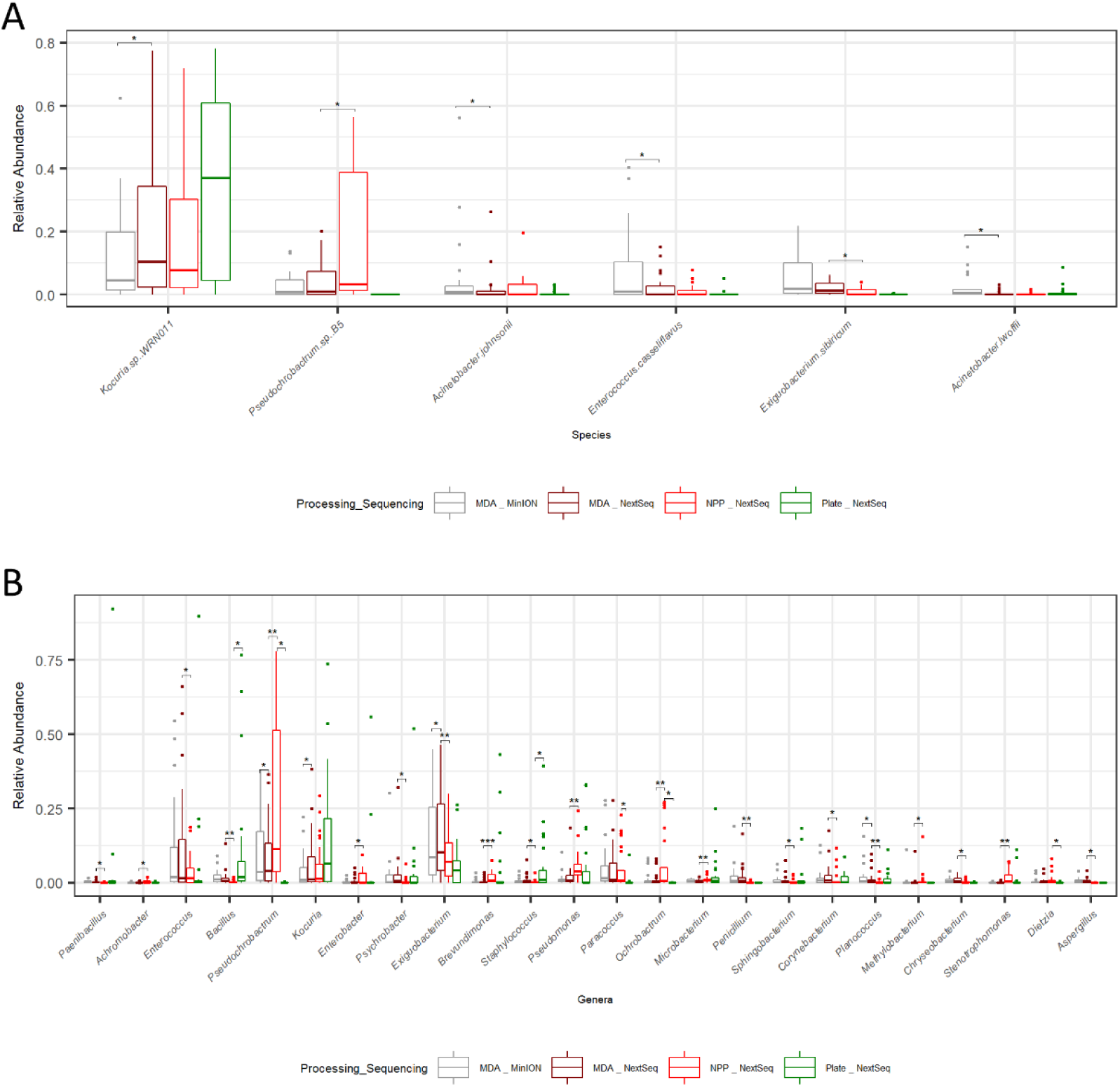
Significant differences in the relative abundance of taxa based on processing and sequencing method. A. Significant species level differences due to sequencing and processing methods on environmental samples. Controls are excluded from these calculations and figures. B. Significant genera level differences in relative abundance due to sequencing and processing methods on environmental samples. Controls are excluded from these calculations and figures.

## Discussion

16S rRNA rapid barcoding-based MinION sequencing of a simple mock community coupled with MEGAN classification by aligning with BLAST against a Silva database provided species level classification to 3 of the 4 species in a mock community and correctly identified both genera present. The rapid sequencing kit-based shotgun sequencing on the MinION platform coupled with LAST alignment against the NR database and MEGAN taxonomic classification resulted in correct classification of all four species but with low level false positive detection of *B. paralicheniformis*, a close relative of *B. licheniformis*. Thus, in this regard, MinION rapid WMGS performed better than MinION 16S sequencing for species level classification of related species, and could be further improved by reducing/eliminating false positives by exercising a stricter cut off and only focusing on species detected at high relative abundance.

Environmental DNA samples subject to MDA resulted in MinION sequencing reads that were shorter, with lower output than the high quality, high quantity, pooled mock metagenomic DNA generated. This is a particular issue for sequencing of low biomass environmental samples where, without the use of MDA, the quantities of DNA would not suffice for current rapid protocols, even after pooling of multiple swabs. The multiplexing of poorer quality DNA from environmental samples resulted in saturation of flow cells, resulting in lower output compared to the mock sequencing run. Despite this, MinION sequencing of environmental samples did perform well and was comparable to other methods when all factors were considered. Some of the most abundant species identified included *Kocuria* sp.

WRN011, *Enterococcus casseliflavus* and *Enterococcus faecium. Kocuria* sp. WRN011 is a saline alkaline soil isolate, and is perhaps selected for due to the unfavourable conditions within a food processing environment arising after cleaning in place (CIP). Both *Enterococcus faecium* and *Enterococcus casseliflavus* are common dairy microorganisms (Rivas *et al*., 2012; Gelsomino *et al*., 2002), with *Enterococcus* sp. been known to also be capable of growth at high pH and in the presence of NaCl (Khedid *et al*., 2009).

However, caution is needed when interpreting the results, particularly from low biomass areas. There did appear to be some cross over between environmental sequences and negative controls particularly in environmental samples with low molecular loads. It must be considered that results for species classified in these samples could be false positives from cross over or contamination of sequences from other samples at any stage of swabbing, extraction, amplification or sequencing. As this occurrence was noted in both MinION and NextSeq generated sequences it is unlikely to be due to barcode misassignment or index swopping alone.

MAG analysis revealed 10 good quality genomes from combined MinION and MDA Illumina sequence reads. Seven of the 10 genomes originated from environmental swab samples, with the other 3 corresponding to positive controls. This form of analysis can, if carried out on a larger scale in the future and with greater sequencing depth, be used to bridge discrepancies in taxonomic classification.

There were also significant differences in the relative abundance of species due to the pre-processing and sequencing approaches taken. MinION sequencing indicated greater relative abundances of *Enterococcus casseliflavus, Acinetobacter lwoffii* and *Acinteobacter johnsonnii* than was suggested by MDA NextSeq sequencing. NextSeq MDA appeared to preferentially sequence *Kocuria sp*. WRN011 compared to MinION. *Pseudochrobactrum sp*. B5 abundances appeared lower in MDA (MinION and NextSeq) and easily culturable NextSeq plate samples than NPP samples. From a culture-based perspective, it is noted that this species is known to reduce hexavalent chromium (Ge, Dong and Zhou, 2013) and it may not grow well on the BHI agar used. *Exiguobacterium sibricum* was detected in higher relative abundance in MDA amplified samples, with significantly higher relative abundance in MDA NextSeq samples compared to NPP NextSeq samples. This suggests it is preferentially amplified by multiple displacement amplification, leading to an overestimation of its relative abundance in these samples.

There were also significant differences in the relative abundances of genera that could not be assigned at the species level. This was most apparent when MDA NextSeq and NPP NextSeq outputs were compared. As well as plate sequences having lower levels of *Pseudochrobactrum*, they were also a lot less diverse than those generated through culture-independent approaches, suggesting culturing at the conditions used was less sensitive. Many species were seen in higher relative abundance in NextSeq plate samples than samples not subject to pre-culturing, including *Planococcus massiliensis, Microbacterium oxydans, Acinetobacter baumannii* and *Lysinibacillus sp*. B2A1, presumably as a consequence of being better suited to growth in these conditions.

Small, portable, real-time DNA sequencers provide the first steps towards real-time industry paced microbial classification and analysis, which could allow the implementation of process change to counteract microbial issues. Although DNA sequencing has been used sporadically for source tracking (Doyle *et al*., 2017; Fretin *et al*., 2018) and monitoring the microbiota through various seasons and environmental conditions (Li *et al*., 2018), there are currently limited numbers of publications and datasets relating to food chain and processing facility microbiomes. While Oxford Nanopore sequencing accuracy is constantly improving (Watson and Warr, 2019), this in itself provides another hurdle to routine implementation in food processing environments, due to often lack of back compatibility with kits, hardware, software and analysis pipelines. More importantly, the need for high quality, high quantity DNA from swabs of an area that actively aims to have low bacterial loads is a challenge, further highlighting the need for adequate controls. Ideally, future forms of portable technologies can be implemented with a rapid kit, without a need for amplification. Despite these challenges, this study and the data generated will aid further attempts to characterise the microbiotas across the food chain, leading to an acceleration towards routine implementation. This is particularly true regarding the generation of MAGs from MDA amplified DNA, resulting in good quality MAGs for 7 environmental isolates, for which relatively few genomes are already available. Notably, in some cases it was difficult to assign some of these MAGs to an existing species, suggesting that the genomes isolated were from related, but previously unclassified species. While *Exiguobacterium sp*. and *Kocuria sp*. have previously been reported in food processing environments (Vishnivetskaya and Kathariou, 2005; Røder *et al*., 2015), *Planococcus sp*., although not well characterised with few genomes available, are regarded as halotolerant, water-associated microorganisms, rather than food processing contaminants (Waghmode *et al*., 2019). The generation of this MAG and further generation of MAGs, will accelerate the identification of food chain microbes through sequencing-based approaches in the future.

Ultimately, while this study highlights issues relating to sourcing sufficient template DNA, inconsistencies across sequencing approaches and platforms, and challenges with assigning taxa, the considerably great potential merits of applying metagenomic approaches to monitor the microbiology of the food chain are clear.

## Materials and methods

### Mock community

DNA from 4 target strains, *Bacillus cereus* DSM 31/ATCC 14579, *Bacillus thuringiensis* DSM 2046/ATCC 10792, *Bacillus licheniformis* DSM 13/ATCC 14580 and *Geobacillus stearothermophilus* DSM 458 (Accession numbers GCF_000007825.1, GCF_002119445.1, GCF_000011645.1 and GCF_002300135.1, respectively), was combined to represent a ‘mock’ metagenomic sample of spore-forming bacteria. Genomic DNA was purchased (latter two strains, DSMZ) or extracted from in-house stocks (former two strains). Where necessary, DNA extraction was performed using the GenElute Bacterial Genomic DNA extraction kit (Sigma Aldrich, NA2110) according to manufacturer’s instructions for Gram positive bacteria DNA extraction except that DNA was eluted in 75 µl elution solution. DNA concentrations were determined using the Qubit double-stranded DNA (dsDNA) high sensitivity (HS) assay kit (BioSciences) and ran on 1% agarose gel to check quality. DNA was diluted to 24 pM and pooled equimolar. 16S rRNA metagenome sequencing, using the 16S rapid barcoding kit SQK-RAB204, as well as rapid whole metagenome sequencing (WMGS), using the rapid sequencing kit SQK-RAD004, was performed using the Oxford Nanopore MinION sequencer. These kits required 10 ng and 400 ng of DNA input, respectively. More specifically, the SQK-RAB204 16S rapid barcoding kit was used for library preparation according to manufacturer’s instructions with barcode 01. DNA was sequenced on FloMIN 106 R9 version flowcell mk1 with minKNOW version 1.7.14 according to manufacturer’s instructions. The SQK-RAD004 rapid sequencing kit was used to prepare the DNA according to manufacturer’s instructions, DNA was sequenced on FloMIN 106 R9 version flowcell mk1 with MinKNOW version 1.11.5 according to manufacturer’s instructions.

### Bioinformatic analysis of mock community metagenomic DNA

Genome sequences for the 4 strains represented in the mock metagenomic community were downloaded from NCBI RefSeq and aligned in a pairwise manner using the Artemis comparison tool (ACT) (Carver *et al*., 2005) (Supplemental Figure 1). 16S DNA sequences were rebasecalled using Albacore (version 2.2.6). FastQC was used to check sequence length and quality. IDBA fq2fa was used to convert fastq files to fasta format. BLASTn alignment (Altschul *et al*., 1990) of sequences against 16S Silva database (release 132) (Pruesse *et al*., 2007; Quast *et al*., 2012) was performed with taxonomic classification by MEGAN (version 6.12.3) (Huson *et al*., 2007). Genus and species levels of classification were determined, and relative abundances calculated and plotted using R ggplot2 (Wickham, 2009). Following basecalling with Albacore, Porechop (version 0.2.4) was used to remove adaptors from rapid WMGS reads before FastQC was used to check sequence length and quality and IDBA fq2fa was used to convert fastq format to fasta format (Peng *et al*., 2012). LAST alignment of reads (Kielbasa *et al*., 2011; Sheetlin *et al*., 2014) was performed against the NR database (March 2018) (Pruitt, Tatusova and Maglott, 2005; Pruitt *et al*., 2012) with MEGAN long read (LR) (MEGAN version 6.12.3) (Huson *et al*., 2018) taxonomic classification. Ranks were split, relative abundances calculated and plotted using R ggplot2 (Wickham, 2009).

The assembly of contigs from metagenomic reads was performed using Canu version 1.7 (Koren *et al*., 2017) with -nanopore-raw flag. MUMmer alignment was performed on the assembled contigs against the 4 known species genomes from RefSeq, with dnadiff used to highlight differences between assemblies and reference genomes (Kurtz *et al*., 2004; Delcher *et al*., 2002). The resulting comparisons were visualised using R ggbio and GenomicRanges (Yin, Cook and Lawrence, 2012; Lawrence *et al*., 2013).

### Environmental sample collection and processing

Environmental swabbing was performed in a commercial dairy processing pilot plant. Eight locations were swabbed during the course of a single day, after cleaning in place (CIP) had been completed and before the next round of dairy processing (Figure 2). These eight locations included a table, door, wall, gaskets/flow plate seals, external surface of dryer balance tank, internal surface of dryer balance tank, external surface of evaporator, and drain beside evaporator. Overall, these eight locations were swabbed over three different months (October, November, December), at a frequency of once per month. Swabbing was performed using Technical Service Consultants Ltd. sponges in neutralising buffer (Sparks Lab Supplies, SWA2023). 5 swabs were performed per surface. Swabbing was performed according to manufacturer’s instructions (Supplemental Methods 1.1). In the laboratory, 5 sponges for each area were pooled aseptically into the stomacher bag of one. Each bag of 5 sponges was subjected to stomaching at 260 rpm for 1 minute. The liquid was then removed, yielding 21 ml for each sample of 5 sponges. 20 ml was prepared for DNA extraction. 1 ml was used for culturing. Two x 15 ml falcon tubes for each sample holding a total of 20 ml were centrifuged at 4,500 x g for 20 min at 4°C. The supernatant was discarded, and pellet resuspended in 500 µl UV treated, autoclaved phosphate buffered saline (PBS). The two resuspended pellets for each area were pooled into a 2 ml microfuge tube. This tube was centrifuged at 13,000 x g for 2 min and the supernatant was discarded. The pellet was stored at -80°C for up to 1 month before DNA extraction. Swab negative controls were also processed in the same way for each sampling day. Briefly, 5 swabs were pooled, subjected to stomaching, liquid collected, 1 ml split for culturing, 20 ml pelleted, washed and frozen.

### Culture analysis

Of the 1 ml of liquid recovered from each stomacher bag, 100 µl was plated on BHI agar in triplicate. Another 100 µl was used for serial tenfold dilution and spread plate on BHI agar in triplicate. All agar plates incubated at 30°C for 48 h. 600 µl of liquid was subjected to spore pasteurisation by heating to 80°C for 12 min in a heating block. This heat treated liquid was then spread plated on BHI in triplicate for incubation at both 30°C and 55°C for 48 h, after which time colonies were counted to determine colony forming units (CFU).

For each sample, the colonies from one agar plate, onto which the neat stomacher bag liquid had been plated, were removed by washing and pelleted to facilitate DNA extraction to represent metagenomic DNA from easy to culture environmental microorganisms. To this end, 5 ml PBS was added to the agar plate, and swirled around, before colonies were scraped off with a sterile Lazy-L spreader (Sigma-Aldrich) and 4 ml recovered into a sterile 15 ml falcon tube. This was centrifuged at 4,500 x g for 20 min at 4°C before removing supernatant. The resulting pellet was resuspended in 1 ml PBS and transferred to a 2 ml microfuge tube. The tube was centrifuged at 13,000 x g for 2 min at room temperature and supernatant removed. The pellet was stored at -80°C for up to three months before DNA extraction. From other agar plates, isolated colonies with obviously different morphologies from each sample were picked, restreaked for purity, inoculated in BHI broth and stocked at -20°C in a final concentration of 25% glycerol.

### DNA extraction and MDA amplification

The Qiagen PowerSoil Pro kit was used for DNA extractions from both environmental sample pellets, and easily culturable washed plate pellets. Easily culturable pellets were removed from -80°C storage and resuspended in 1 ml PBS. 200 µl (or 500 µl for 9 smaller pellets, corresponding door, external evaporator and internal dryer balance tank samples for all 3 months) was removed and centrifuged at 12,000 x g for 2 min. The supernatant was discarded and the pellet retained. These pellets, and those sourced directly from environmental swabbing, i.e. without culture, were resuspended in 800 µl CD1 and transferred to a Powerbead Pro tube. Powerbead Pro tubes were secured in a tissue lyser set at 20 Hz for 10 min before centrifuging and following the rest of the PowerSoil Pro kit manufacturer’s instructions, eluting in a smaller volume, of 35 µl. For each sampling day, negative controls, involving unused swabs, were also prepared by following an identical extraction protocol and additional negative controls, to detect kit contaminants, were generated whereby an extraction was performed using the kit reagents alone, starting with 800 µL solution CD1.

Whole metagenome amplification was performed using multiple displacement amplification (MDA) with the REPLI-g Single Cell kit (Qiagen, 150345). MDA was performed using DNA from environmental samples and controls for each day. These controls consisted of swab negative control, DNA extraction kit negative control, blank MDA preparation as a MDA negative control and mock metagenomic community (section 1.6.1) as a positive control. DNA concentrations were determined using Qubit dsDNA HS kit. Samples with high DNA concentrations were diluted such that all samples had a final concentration of < 10 ng in 2.5 µl. MDA amplification was performed according to manufacturer’s instructions (Supplemental Methods 1.2) for 12 sample amplifications at a time (8 environmental samples, 1 positive control, 3 negative controls (swab, extraction, MDA)). Amplified DNA was then stored at -20°C.

### Library preparation and sequencing

#### MinION library

DNA concentrations of 36 MDA samples were measured using both the Qubit dsDNA broad range (BR) and HS assays and diluted to 400 ng in 7.5 µl. Three libraries were prepared, containing 12 samples each (8 environmental MDA samples, 3 MDA negative controls (swab, extraction and MDA kit negative controls) and a MDA mock community positive control) per flow cell. The SQK-RBK004 rapid barcoding kit was used to prepare the DNA according to manufacturer’s instructions, including an optional Ampure XP clean up step, directly prior to sequencing. DNA was sequenced on FloMIN 106 R9 version flowcell mk1 with MinKNOW version 18.12.4 according to manufacturer’s instructions.

### Illumina Nextera library

The DNA concentrations of MDA (n = 36), non-MDA (i.e., metagenomic DNA not subjected to pre-processing (NPP)) (n = 33), and easily culturable (Plate) (n = 24) metagenomic DNA samples was measured using the Qubit HS dsDNA kit and diluted. DNA was prepared for Illumina sequencing following Illumina Nextera XT Library Preparation Kit guidelines except that tagmentation was performed for 7 min. DNA tagmentation was visualised using Agilent Bioanalyzer high sensitivity DNA analysis, and average fragment size calculated. The DNA concentration was measured by Qubit HS dsDNA assay and the concentration then calculated, before diluting and pooling at equimolar ratios. The DNA library was sequenced on Illumina NextSeq at the Teagasc DNA sequencing facility, with a NextSeq (500/500) High Output 300 cycles v2.5 kit (Illumina 20024908).

### 16S rDNA Sanger sequencing of isolated colonies

16S colony PCR was performed (Supplemental Methods 1.3) using universal primers 27F and 338R for 16S gene (AGAGTTTGATCCTGGCTCAG and CATGCTGCCTCCCGTAGGAGT, respectively). PCR products were run on a 1 % agarose gel, before cleaning with 1.8 x Ampure XP. 5 µl of each cleaned up PCR product was aliquoted into a 96 well plate and 5 µl of forward primer added on top at 5 µM according to GATC requirements. A unique barcode was added to each plate and sent to GATC Biotech (Germany) for Sanger sequencing. A subset of amplicons were also sequenced with the reverse primer to ensure accuracy.

### Bioinformatic analysis of environmental metagenomic DNA Analysis of MinION data

Guppy basecalled reads obtained from MinKnow (version 18.12.4) were demultiplexed using Guppy barcoder version (2.1.3) to produce a barcoding summary text file. This contained the percentage match of each read to their barcodes with a minimum score of 60, the default). All fastq files produced by MinKnow were concatenated and guppy_bcsplit.py (https://github.com/ms-gx/guppy_bcsplit) allowed demultiplexing of reads based on their barcode assigned in the barcoding summary text file. Porechop (version 0.2.4) was used to remove adaptors from rapid kit sequence reads before Fastqc was used to check sequence length and quality. IDBA fq2fa was used to convert fastq to fasta (Peng *et al*., 2012). LAST alignment of fasta files (Kielbasa *et al*., 2011; Sheetlin *et al*., 2014) against the NR database (March 2018) (Pruitt, Tatusova and Maglott, 2005; Pruitt *et al*., 2012) was performed with the MEGAN LR classification (MEGAN version 6.12.3) (Huson *et al*., 2018). Files were merged, ranks were split, total number of bases sequenced, and classified were calculated. Relative abundances calculated and plotted using R ggplot2 (Wickham, 2009).

### Analysis of NextSeq data

BCl2fastq was used to convert raw sequence reads from Illumina NextSeq to fastq format. Kneaddata from bioBakery (McIver *et al*., 2018) used trimmomatic for quality filtering and trimming paired end files (Bolger, Lohse and Usadel, 2014) with BMTagger to remove human and bovine reads. FastQC was used to visualise sequence length and quality. IDBA converted fastq to fasta (Peng *et al*., 2012).

Diamond alignment (Buchfink, Xie and Huson, 2015) of fasta files was performed against the NR database (march 2018) (Pruitt, Tatusova and Maglott, 2005; Pruitt *et al*., 2012) with MEGAN classification (MEGAN version 6.12.3) (Huson *et al*., 2018). Files were merged, ranks were split, total number of bases sequenced and classified calculated and relative abundances calculated and plotted using R ggplot2 (Wickham, 2009).

Illumina data was also analysed using Kraken2 and Bracken (Lu *et al*., 2017; Wood and Salzberg, 2014) for taxonomy classification as well as using MetaPhlan2 (Truong *et al*., 2015) for taxonomy classification for the purpose of comparison.

### Generation of MinION-Illumina hybrid Metagenome-assembled genomes

MDA amplified sequences from both Illumina and Oxford Nanopore sequencing were assembled using OPERA-MS (Bertrand *et al*., 2019). Illumina reads were then mapped against assemblies using bowtie2 (Langmead and Salzberg, 2012) and bam files sorted using samtools (Li *et al*., 2009). Depth was calculated and Metabat2 ran on assembled contigs to produce bins (Kang *et al*., 2015; Kang *et al*., 2019). Checkm was used to determine the quality of the metagenome assembled genomes (MAGs). Prokka (Seemann, 2014) was used to generate .ffn files from bins, Kaiju (Menzel, Ng and Krogh, 2016)-based taxonomic classification was performed on the open reading frames from prokka. Megan LR (Huson *et al*., 2018) was also used on the whole bins for taxonomic classification of high quality MAGs.

### Culture-and 16S rRNA Sanger sequence-based analysis

CFUs were determined on the basis of an average of three agar plates per sample. CFU per swab was calculated by dividing by 5 (5 swabs=1 sample, and each swab covered area 360cm^2^). 16S rRNA Sanger sequences resulting from morphologically different isolates per sample were blasted using BLASTn against the 16S ribosomal RNA (Bacteria and Archaea) database on NCBI, with top hits recorded, and genus level classification analysed.

### Statistics

Pairwise Wilcoxon rank sums test using Benjamini Hochberg *p*-value correction analysis was used to compare sample groups, including investigations of the impact of sequencer type on taxonomy classification with MinION MDA-treated and NextSeq MDA-treated samples. The impact of MDA amplification was also investigated in this way through comparison between NextSeq MDA treated samples and NextSeq no pre-processing (NPP) samples. Differences in taxonomy classification between sequences derived from environmental metagenomic DNA versus those sourced from easy to culture microorganisms was shown by comparing NextSeq NPP and NextSeq easy to culture (plate) sequences. Diversity analysis was performed in R with vegan package. Shannon and Simpson alpha diversity metrics were calculated along with Bray Curtis Nonmetric Multidimensional Scaling beta diversity metrics. Pairwise Wilcoxon rank sums test using Benjamini Hochberg p-value correction was used to compared samples groups based on sequencing and processing methods used, controls were excluded from these calculations.

## Accession number

Sequence data have been deposited in the European Nucleotide Archive (ENA) under the study accession number PRJEB39267.

## Acknowldegements

This research was funded by the Department of Agriculture, Food and the Marine (DAFM), under the FIRM project SACCP, reference number 14/F/883. Research in the Cotter laboratory is also funded by Science Foundation Ireland (SFI) under grant numbers SFI/12/RC/2273 (APC Microbiome Ireland) and SFI/16/RC/3835 (Vistamilk) and by the European Commission under the Horizon 2020 program under grant number 818368 (Master).

