## Supplemental documents for "Microbiome-based environmental monitoring of a dairy processing facility highlights the challenges associated with low microbial-load samples"

### Supplemental Figures

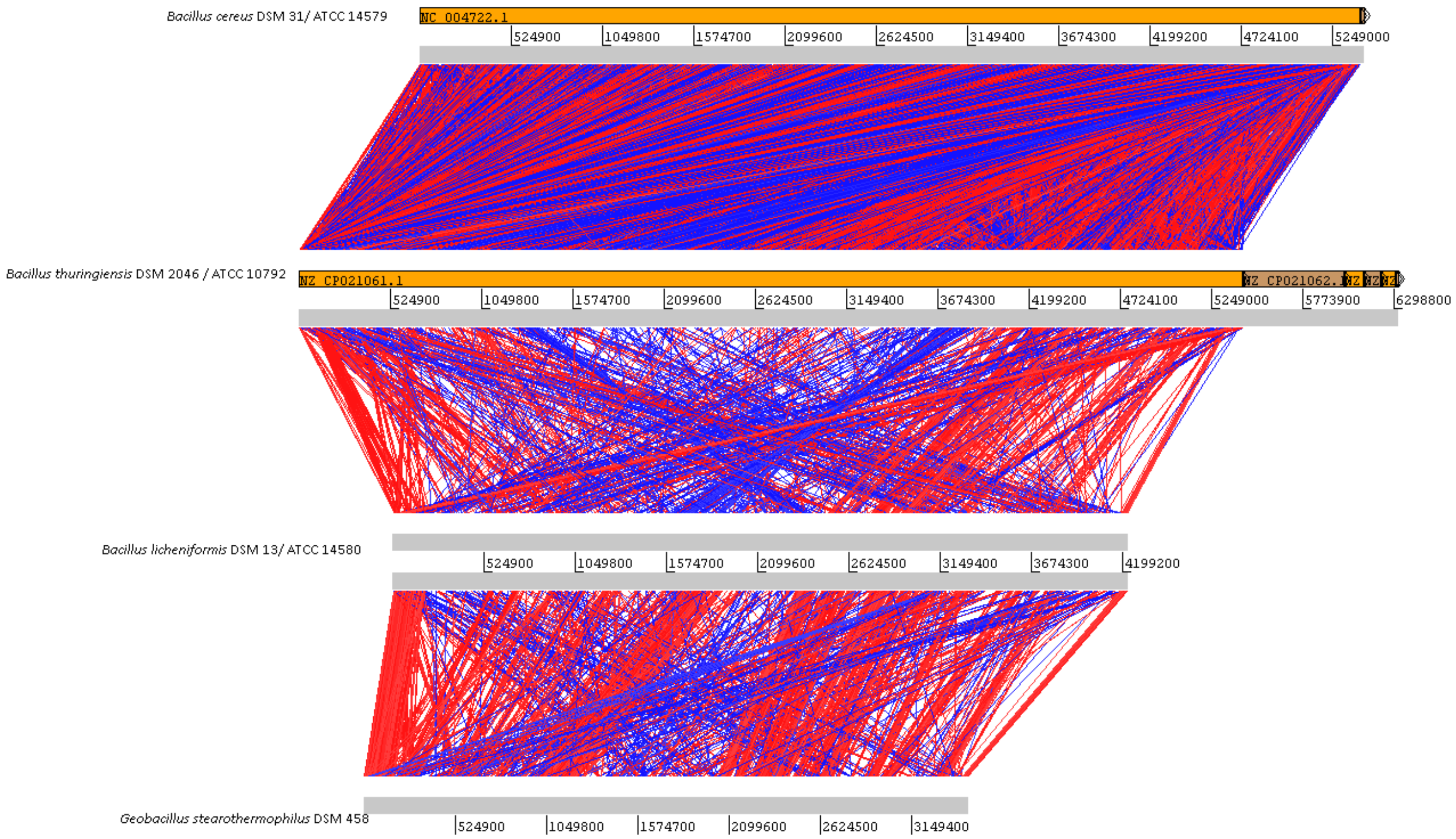


Supplemental Figure 1. Comparison of genomes of 4 strains used in mock community using ACT.

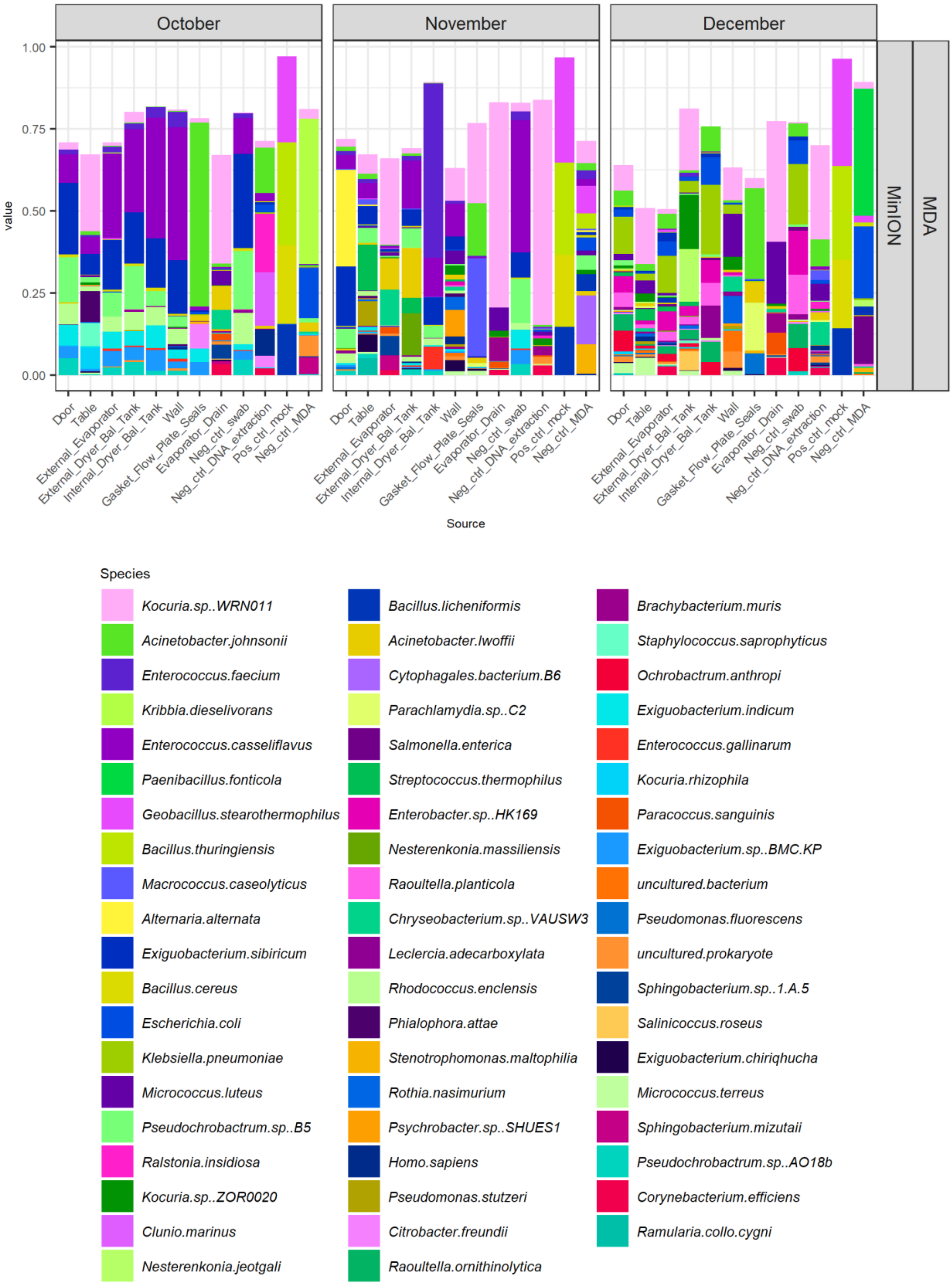


Supplemental Figure 2. Species level classification of MinION environmental samples.

Oxford Nanopore Technologies MinION sequencing of MDA DNA from environmental swab samples classified using LAST+MEGAN LR. Species present > 5% relative abundance in at least one sample are shown.

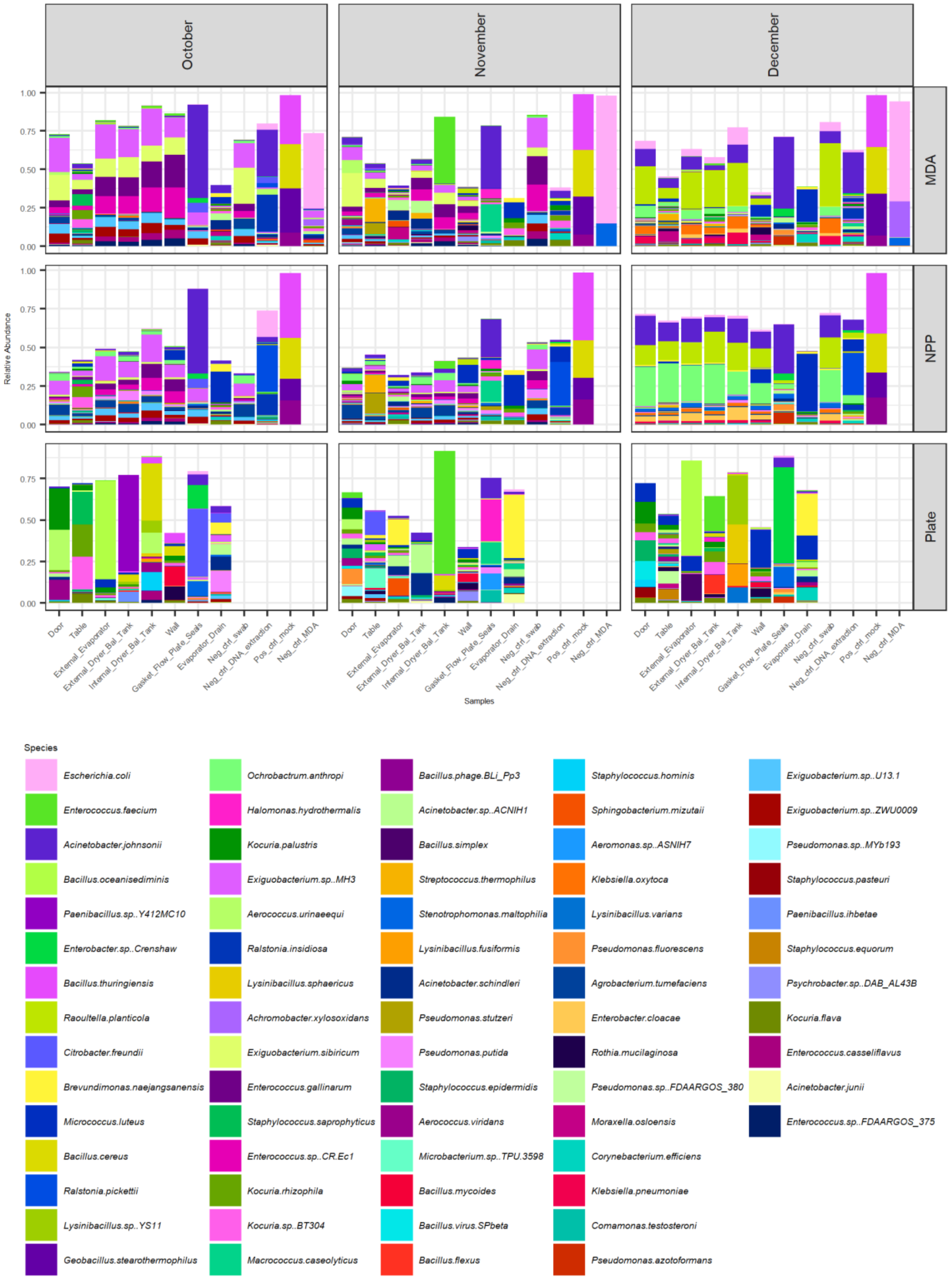


Supplemental Figure 3. Bracken classification of NextSeq sequences.

Kraken2 with Bracken species level classification of NextSeq samples. 100% mock community was classified, but yet could not accurately decipher species level classification.

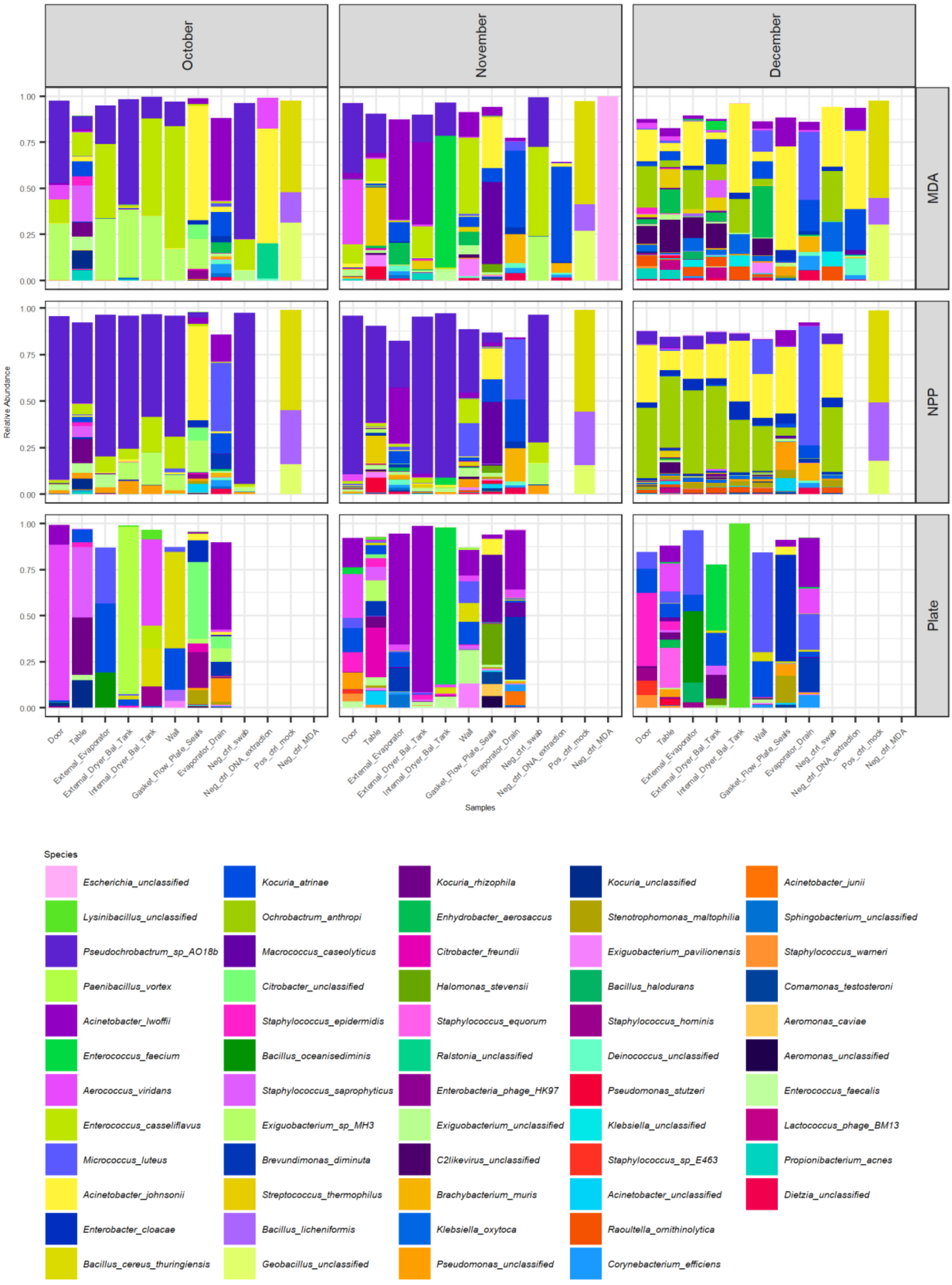


Supplemental Figure 4. MetaPhlAn2 species level classification of NextSeq data. MetaPhlAn2 species level classification of NextSeq data. Again highlights an incorrectly classified positive control/mock community.

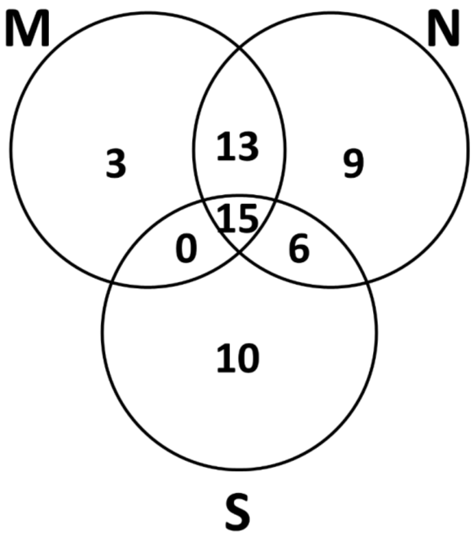


Supplemental Figure 5. Venn diagram of the number of genera assigned per sequence type.

Venn diagram shows number of genera assigned per sequencing type, MinION > 5% relative abundance (M), NextSeq > 5% relative abundance (N), Sanger (S).

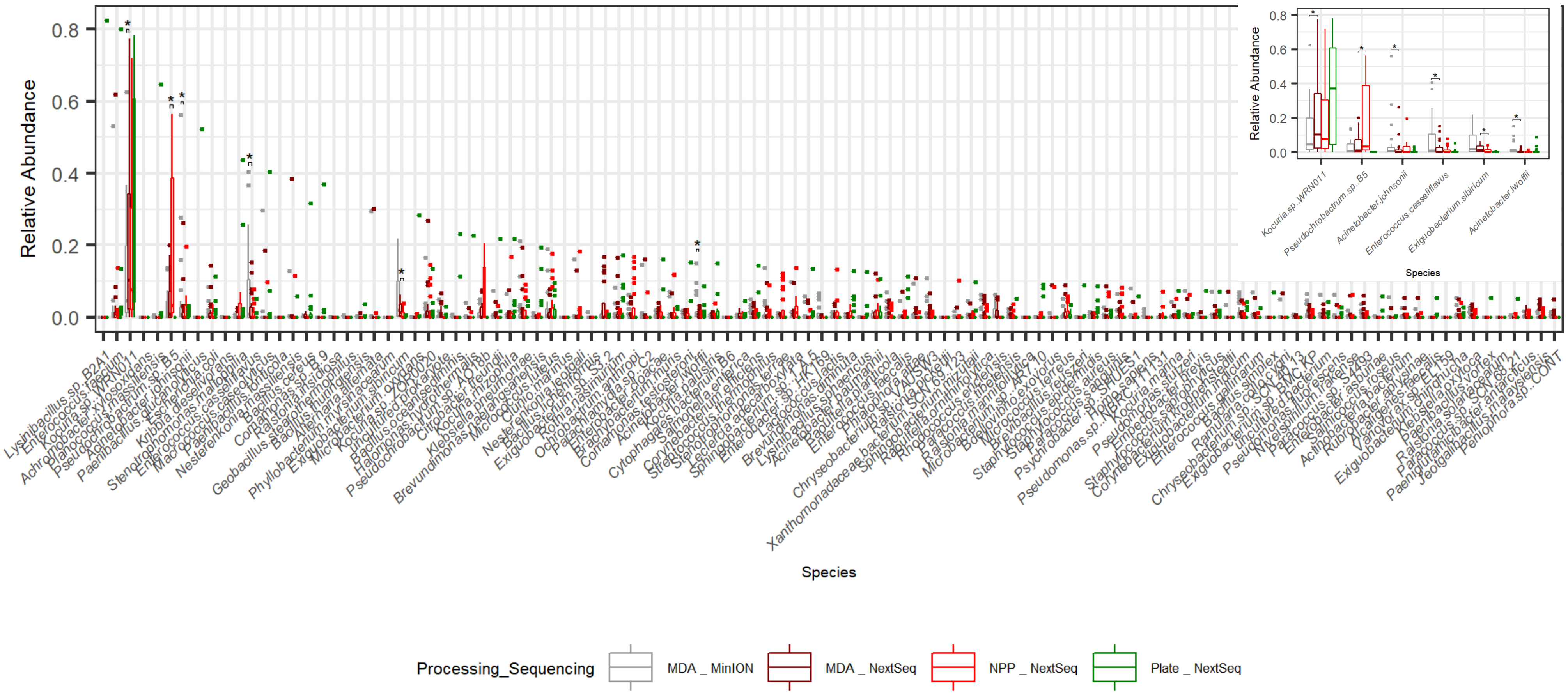

37

38 Supplemental Figure 6. Distribution of species level classification in environmental samples

Distribution of species classification by sequencer and pre-processing type. Significant differences highlighted. Controls not included in these, only environmental samples.

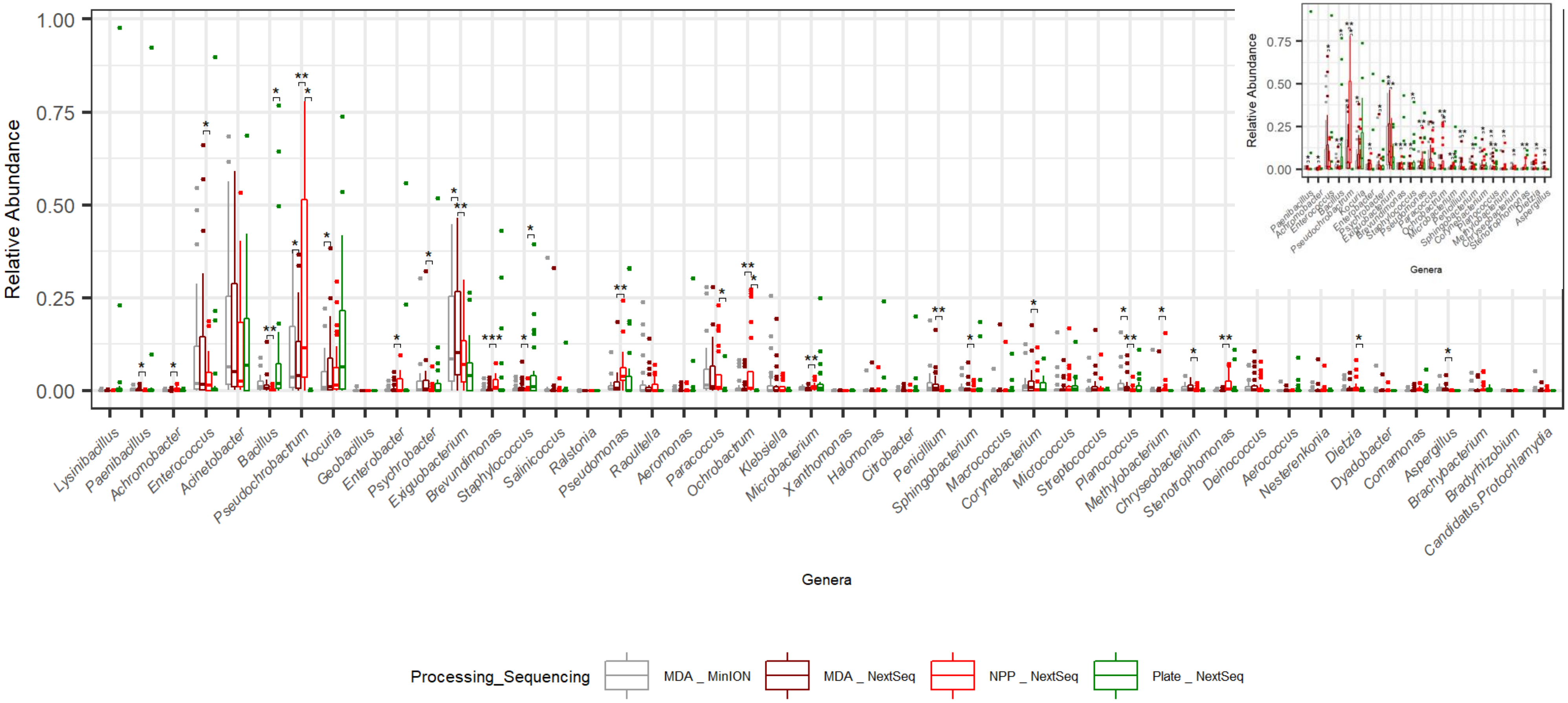


Supplemental Figure 7. Distribution of genera level classification in environmental samples.

Distribution of genera classification by sequencer and pre-processing type. Significant differences highlighted. Controls not included in these, only environmental samples.

**Supplemental Tables**

Supplemental Table 1. Alignment of sequenced mock metagenome assembled genomes (QRY) to reference genomes from NCBI RefSeq (REF), using MUMmer.

| | [REF] | [QRY] |
|---|---|---|
| [Sequences] | | |
| TotalSeqs | 10 | 104 |
| AlignedSeqs | 9(90.00%) | 104(100.00%) |
| UnalignedSeqs | 1(10.00%) | 0(0.00%) |
| [Bases] | | |
| TotalBases | 19432784 | 17433642 |
| AlignedBases | 19095736(98.27%) | 17362744(99.59%) |
| UnalignedBases | 337048(1.73%) | 70898(0.41%) |
| [Alignments] | | |
| 1-to-1 | 243 | 243 |
| TotalLength | 17330360 | 17043215 |
| AvgLength | 71318.35 | 70136.69 |
| AvgIdentity | 97.94 | 97.94 |

Supplemental Table 2. Counts of culturable bacteria from environmental samples, and their classification.

| Source | Month | Culture type | CFU / Sample | CFU / Swab | Genera |
|---|---|---|---|---|---|
| Door | October | TBC | 2.00E+04 | 4.00E+03 | *Acinetobacter, Aerococcus, Aerococcus, Kocuria, Kocuria, Psychrobacter* |
| | | ST | 0.00E+00 | 0.00E+00 | N/A |
| | | SM | 0.00E+00 | 0.00E+00 | N/A |
| Table | October | TBC | 3.34E+05 | 6.69E+04 | *Aerococcus, Bacillus, Kocuria* |
| | | ST | 7.00E+01 | 1.40E+01 | *Bacillus* |
| | | SM | 1.05E+04 | 2.10E+03 | *Bacillus, Bacillus, Bacillus, Bacillus, Paenibacillus, Paenibacillus* |
| External Evaporator | October | TBC | 5.88E+03 | 1.18E+03 | *Bacillus, Lysinibacillus, Micrococcus* |
| | | ST | 3.50E+02 | 7.00E+01 | *Bacillus* |
| | | SM | 2.94E+03 | 5.88E+02 | *Bacillus, Bacillus, Bacillus* |
| External Dryer Bal Tank | October | TBC | 9.38E+03 | 1.88E+03 | *Bacillus, Bacillus, Bacillus, Chryseomicrobium, Exiguobacterium, Lysinibacillus* |
| | | ST | 7.00E+01 | 1.40E+01 | N/A |
| | | SM | 7.00E+01 | 1.40E+01 | N/A |
| Internal Dryer Bal Tank | October | TBC | 3.22E+03 | 6.44E+02 | *Bacillus, Lysinibacillus, Staphylococcus, Staphylococcus* |
| | | ST | 0.00E+00 | 0.00E+00 | N/A |
| | | SM | 0.00E+00 | 0.00E+00 | N/A |
| Wall | October | TBC | 9.87E+03 | 1.97E+03 | *Bacillus, Bacillus, Microbacterium, Rothia, Rothia* |
| | | ST | 7.70E+02 | 1.54E+02 | *Bacillus* |
| | | SM | 1.82E+03 | 3.64E+02 | *Oceanobacillus* |
| Gasket Flow Plate Seals | October | TBC | 3.86E+08 | 7.71E+07 | *Acinetobacter* |
| | | ST | 0.00E+00 | 0.00E+00 | N/A |
| | | SM | 7.06E+04 | 1.41E+04 | *Bacillus, Lysinibacillus* |
| Evaporator Drain | October | TBC | 2.85E+08 | 5.70E+07 | *Corynebacterium, Kocuria, Psychrobacter* |
| | | ST | 5.25E+02 | 1.05E+02 | *Bacillus* |
| | | SM | 7.50E+04 | 1.50E+04 | N/A |
| Neg ctrl Swab | October | TBC | 7.00E+01 | 1.40E+01 | *Staphylococcus* |
| | | ST | 0.00E+00 | 0.00E+00 | N/A |
| | | SM | 0.00E+00 | 0.00E+00 | N/A |
| Door | November | TBC | 1.90E+04 | 3.81E+03 | *Aerococcus, Bacillus, Kocuria, Staphylococcus, Staphylococcus, Staphylococcus, Staphylococcus, Staphylococcus* |
| | | ST | 3.15E+02 | 6.30E+01 | *Bacillus* |

| | | | | | |
|---|---|---|---|---|---|
| | | SM | 3.15E+02 | 6.30E+01 | *Bacillus* |
| Table | November | TBC | 1.53E+05 | 3.06E+04 | *Bacillus, Bacillus, Exiguobacterium, Exiguobacterium, Kocuria, Microbacterium* |
| | | ST | 2.80E+02 | 5.60E+01 | N/A |
| | | SM | 1.05E+03 | 2.10E+02 | *Bacillus, Lysinibacillus, Rothia* |
| External Evaporator | November | TBC | 1.24E+07 | 2.48E+06 | *Acinetobacter, Kocuria, Microbacterium, Microbacterium, Sphingobacterium* |
| | | ST | 7.00E+01 | 1.40E+01 | *Bacillus* |
| | | SM | 3.57E+03 | 7.14E+02 | *Bacillus, Bacillus, Bacillus, Kocuria* |
| External Dryer Bal Tank | November | TBC | 9.28E+04 | 1.86E+04 | *Acinetobacter, Bacillus, Bacillus, Chryseomicrobium, Paracoccus* |
| | | ST | 7.00E+01 | 1.40E+01 | *Bacillus* |
| | | SM | 2.00E+03 | 3.99E+02 | *Bacillus, Bacillus, Bacillus* |
| Internal Dryer Bal Tank | November | TBC | 1.73E+04 | 3.46E+03 | *Enterococcus, Enterococcus, Lysinibacillus* |
| | | ST | 0.00E+00 | 0.00E+00 | N/A |
| | | SM | 1.26E+03 | 2.52E+02 | *Bacillus* |
| Wall | November | TBC | 2.18E+05 | 4.37E+04 | *Kocuria, Kocuria, Lysinibacillus, Planococcus, Psychrobacter, Rothia, Rothia* |
| | | ST | 4.20E+02 | 8.40E+01 | *Bacillus* |
| | | SM | 7.98E+03 | 1.60E+03 | *Bacillus, Bacillus, Bacillus* |
| Gasket Flow Plate Seals | November | TBC | 2.12E+07 | 4.24E+06 | *Acinetobacter, Halomonas, Kocuria, Macrococcus* |
| | | ST | 7.00E+01 | 1.40E+01 | *Bacillus* |
| | | SM | 5.85E+04 | 1.17E+04 | *Bacillus, Lysinibacillus, Lysinibacillus* |
| Evaporator Drain | November | TBC | 2.15E+09 | 4.30E+08 | *Kocuria, Microbacterium, Planococcus* |
| | | ST | 1.47E+03 | 2.94E+02 | *Bacillus, Bacillus* |
| | | SM | 3.53E+05 | 7.06E+04 | *Bacillus, Bacillus, Corynebacterium* |
| Neg ctrl swab | November | TBC | 2.10E+02 | 4.20E+01 | *Bacillus, Kocuria, Rothia* |
| | | ST | 0.00E+00 | 0.00E+00 | N/A |
| | | SM | 2.10E+02 | 4.20E+01 | *Kocuria* |
| Door | December | TBC | 1.61E+04 | 3.22E+03 | *Arthrobacter, Micrococcus, Psychrobacillus, Psychrobacillus, Staphylococcus, Staphylococcus* |
| | | ST | 7.00E+01 | 1.40E+01 | *Bacillus* |
| | | SM | 1.40E+02 | 2.80E+01 | N/A |
| Table | December | TBC | 2.10E+05 | 4.19E+04 | *Bacillus, Kocuria, Rothia, Macrococcus, Rothia, Rothia, Staphylococcus* |
| | | ST | 6.30E+02 | 1.26E+02 | *Bacillus* |
| | | SM | 8.40E+02 | 1.68E+02 | N/A |

| | | | | | |
|---|---|---|---|---|---|
| External Evaporator | December | TBC | 7.56E+03 | 1.51E+03 | *Bacillus, Ornithinibacillus, Ornithinibacillus* |
| | | ST | 1.26E+03 | 2.52E+02 | *Bacillus, Brevibacillus* |
| | | SM | 3.08E+03 | 6.16E+02 | *Bacillus, Bacillus* |
| External Dryer Bal Tank | December | TBC | 4.92E+04 | 9.84E+03 | *Bacillus, Kocuria, Marinilactibacillus, Rothia, Salinicoccus* |
| | | ST | 3.15E+02 | 6.30E+01 | *Bacillus* |
| | | SM | 3.50E+02 | 7.00E+01 | N/A |
| Internal Dryer Bal Tank | December | TBC | 7.00E+01 | 1.40E+01 | *Lysinibacillus, Lysinibacillus* |
| | | ST | 0.00E+00 | 0.00E+00 | N/A |
| | | SM | 7.00E+01 | 1.40E+01 | *Bacillus* |
| Wall | December | TBC | 2.17E+05 | 4.33E+04 | *Aerococcus, Bacillus, Kocuria, Planococcus, Rothia* |
| | | ST | 1.05E+02 | 2.10E+01 | *Thermoactinomyces* |
| | | SM | 1.47E+03 | 2.94E+02 | *Oceanobacillus* |
| Gasket Flow Plate Seals | December | TBC | 7.06E+07 | 1.41E+07 | *Acinetobacter, Acinetobacter, Chryseobacterium* |
| | | ST | 8.40E+02 | 1.68E+02 | *Bacillus* |
| | | SM | 4.57E+04 | 9.14E+03 | *Bacillus, Bacillus, Exiguobacterium* |
| Evaporator Drain | December | TBC | 2.94E+08 | 5.88E+07 | *Aerococcus, Brevundimonas, Sphingobacterium* |
| | | ST | 8.72E+03 | 1.74E+03 | *Brevibacillus* |
| | | SM | 1.17E+05 | 2.35E+04 | *Bacillus, Bacillus* |
| Neg ctrl swab | December | TBC | 1.40E+02 | 2.80E+01 | *Bacillus* |
| | | ST | 0.00E+00 | 0.00E+00 | N/A |
| | | SM | 7.00E+01 | 1.40E+01 | *Bacillus* |

Counts and classification of BHI cultured samples (TBC) and mesophilic and thermophilic spore selected BHI cultured samples (SM and ST) following isolation of morphologically different isolates and identification by sanger sequencing of 16S region.

**Supplemental Documents**

Supplemental Methods 1.1

Hygiene sponge sampling kits swabbing procedure

Briefly, a stomacher bag containing a pre-moistened sponge was shaken to bring the sponge to the bottom of the bag. The bag was torn open above the zip lock; then, holding the sponge at the bottom from the outside, the bag was carefully peeled back, from above the zip lock over the gloved hand, taking care not to touch the inside of the bag or sponge. The exposed sponge was the used to swab an area of 360 $cm^2$ swabbing vertically with one side of the sponge and horizontally over the same area with the other side of the sponge. The bag was then carefully reverted to its original position, without touching the inside and the zip lock sealed. 5 swabs of each area to be sampled were performed in this way. The surface area was then wiped with disinfectant to remove neutralising buffer.

Supplemental Methods 1.2

MDA amplification

Briefly, 500 µl of $H_2O$ was added to buffer DLB, mixed well and centrifuged briefly, storing at -20°C for up to 6 months. Buffer D1 and N1 were prepared according to instructions on the day of use. 2.5 µl buffer D1 was added to 2.5 µl DNA, this was mixed by vortexing and centrifuged briefly. Samples were incubated at room temperature for 3 min. 5 µl buffer N1 was added, mixed by vortexing and centrifuged briefly before storing on ice. Mastermix was prepared on ice according to manufacturer's instructions. For each amplification 40 µl master mix was added to 10 µl denatured DNA. This was incubated on thermocycler at 30°C for 8 h.

The polymerase was then inactivated by heating samples to 65°C for 3 min also in the thermocycler. Thermocycler settings ( 2 x 4 h holds at 30°C and 1 x 3 min hold at 65°C).

Supplemental Methods 1.3

16S colony PCR

Colonies were picked, and mixed in 50 µl PCR water, before microwaving on full power for 1 minute to disrupt cells.  Master mix consisting of 5 µl AccuTaq LA 10x buffer, 2.5 µl 10 mM dNTP mix, 1 µl DMSO, 2 µl 10 µM Forward primer, 2 µl 10 µM reverse primer, 32 µl PCR water, 0.5 µl AccuTaq LA DNA polymerase (Sigma Aldrich, D8045) per amplification was made. 45 µl mastermix was added to each tube of 5 µl disrupted colony, before centrifuging briefly to mix and placing on pre-programmed thermocycler with 95°C x 5 min, 25 cycles of 95°C x 30 sec, 55°C x 30 sec, 72°C x 30 sec and a final 72°C x 5 min, before holding at 4°C.
